# HySyn: A genetically-encoded synthetic synapse to rewire neural circuits *in vivo*

**DOI:** 10.1101/2020.01.27.922203

**Authors:** Josh D. Hawk, Daniel A. Colón-Ramos

## Abstract

Here we introduce HySyn, a system designed to rewire neural connectivity *in vivo* by reconstituting a functional heterologous synapse. We demonstrate that genetically targeted expression of the two HySyn components, a *Hydra*-derived neuropeptide and its receptor, creates *de novo* neuromodulatory transmission in a mammalian neuronal tissue culture model and rewires a behavioral circuit *in vivo* in the nematode *Caenorhabditis elegans*. HySyn can interface with existing optogenetic, chemogenetic and pharmacological approaches to functionally probe synaptic transmission, dissect neuropeptide signaling, or modulate specific neural circuits.

---

Recent advances in optogenetic and chemogenetic tools have allowed unprecedented *in vivo* access to neural circuits^1,2^. While these powerful tools provide specific control over neuronal activity, a gap exists in our ability to similarly manipulate neuronal synaptic communication. Specifically, there is no existing technology to create new, synthetic and manipulatable chemical synaptic connections that enable re-configuring of neural networks in organisms. The ability to control intercellular signaling through creation of synthetic chemical synapses at will could be transformative for efforts aimed at understanding neuromodulation of synaptic connections and dissection of circuit logic.

For this reason, we developed a system to engineer synthetic synapses and elicit control over neuromodulatory connectivity^3,4^. We prioritized design of a system that would be 1) versatile to function in a wide-range of cell-types and organisms, 2) modular to allow independent genetic targeting of pre- and post-synaptic components, 3) specific to modulate only the intended target cells while being inert to endogenous neurotransmission, 4) robust by targeting conserved intracellular signaling cascades, and 5) synergistic to interface with existing optogenetic and chemogenetic technologies. Informed by these goals, we engineered ‘HySyn’, a two-component system that creates a synthetic neuromodulatory synapse to manipulate intracellular calcium within *in vivo* neural circuits.

HySyn is composed of a presynaptic carrier molecule for heterologous production of a cnidarian neuropeptide and a postsynaptic cognate receptor that fluxes calcium^5,6^. The primary sequence of this *Hydra*-derived peptide is distinct from RFamide-related peptides found in other organisms^7,8^ and specific to its receptor^5^. We reasoned that the divergent evolution of this neuropeptide-receptor pair could be exploited to produce a synthetic synapse that would be inert to endogenous neuropeptides unless the intended ligand is also expressed. We also reasoned that because the neuropeptide processing, transport and release mechanisms^12^ and calcium signaling are conserved throughout evolution^9^, expression of this neuropeptide ligand-receptor pair in heterologous systems would enable neuromodulatory control of a fundamental and conserved signal, intracellular calcium^9–11^ at nanomolar neuropeptide concentrations^5^.

To drive heterologous expression, processing, and transport of the cnidarian neuropeptide, we designed a genetically encoded pre-pro-peptide carrier, ‘HyPep’, that harnesses the universality of the neuropeptide processing pathway (Fig S1 and Fig 1a, ‘*Hydra* RFamide). Previous approaches to label neuropeptides concatenated a reporter onto an existing full-length natural neuropeptide precursor^12,13^. We built upon the knowledge of neuropeptide synthesis from these and other studies^8,12–14^ to create HyPep as a novel synthetic pre-pro-peptide carrier that would enable targeting and processing of heterologous neuropeptides using the endogenous neuropeptide processing pathway (Fig S1a). HyPep consists of a signal peptide that directs trafficking, acidic spacers with enzymatic recognition sites for cleavage of the neuropeptide, and the sequence encoding the heterologous neuropeptide itself. We based our signal peptide on neuropeptide Y (Fig S1b, ‘Signal Peptide’), a ubiquitously expressed neuropeptide in vertebrates^12^. We designed artificial neuropeptide spacers containing consensus cleavage sites (Fig 1b, red lines) for pre-pro-convertase (PC2), a conserved neuropeptide endopeptidase^13^. Because cross-species alignment of neuropeptides revealed a strong bias for acidic residues between the dibasic cleavage site, we created acidic linkers between these cleavage sites, which in turn flanked the heterologous cnidarian neuropeptide we sought to express with the HyPep system (Fig 1b, ‘Hydra RFamide’).

**Figure 1.**
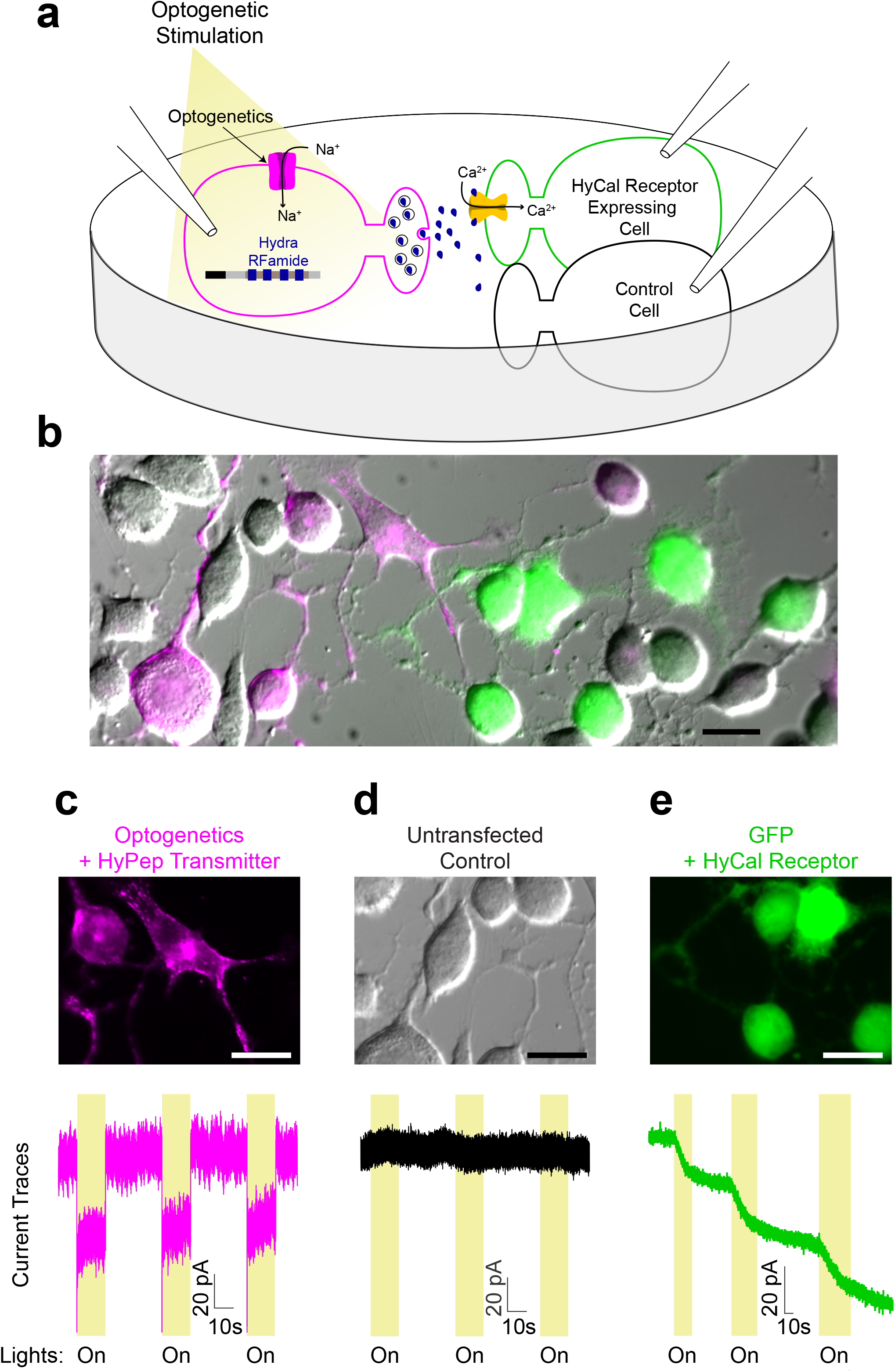
*Hydra*-derived synthetic synapse (HySyn) engineered through heterologous expression of *Hydra* neuropeptide (HyPep) and receptor (HyCal). **a**, Schematic illustrating the experimental paradigm to assay synthetic synapse function by whole-cell patch-clamp electrophysiology. **b**, Micrograph showing co-culture of Neuro2a neuroblastoma cells expressing either presynaptic HyPep with ChRoME (magenta), postsynaptic HyCal with GFP (green) or untransfected (grayscale). **c-e**, Identification of cell populations based on fluorescent markers (top, micrograph, 20μm scale bar) enabling whole-cell current recording (bottom, ‘current traces’) from three distinct populations during optogenetic stimulation (480nm, yellow-shaded windows, ‘On’): (**c**) magenta presynaptic cells expressing HyPep with ChRoME consistently produced step-wise optogenetic currents, (**d**) unlabeled cells remain unaltered by optogenetic activation of neighboring HyPep+ cells, and (**e**) green postsynaptic cells expressing the HyCal receptor show an increasing inward current during light stimulation. Downward deflection of traces during optogenetic stimulation in yellow windows indicates an inward current, suggesting a depolarizing cation current. Persistences of HyCal current between stimulation (**e**) is consistent with the lack of desensitization of this channel and suggests an accumulation of HyPep-derived neuropeptide in bath solution.

Expression of the HyPep synthetic pre-pro-peptide carrier in mammalian Neuro2A cells resulted in localization and transport of a fluorescent reporter to the expected intracellular compartments and vesicular release sites (Fig S1). This observation is consistent with our hypothesis that the engineered HyPep carrier harnesses the universality of the neuropeptide processing pathway to target the *Hydra* neuropeptide processing, transport and release. Next, we used whole-cell patch-clamp electrophysiology to test whether the presynaptic HyPep, when paired with the postsynaptic HyCal receptor, is capable of producing a functional neuromodulatory connection (Fig 1a). We created an optogenetically excitable population of ‘presynaptic’ cells by co-expressing HyPep with mRuby-labeled ChRoME^15^, an optogenetic tool for depolarizing neurons (Fig 1b, magenta). We then co-cultured these presynaptic cells with other cells expressing the ‘postsynaptic’ receptor HyCal (Fig 1b, green), as well as untransfected control cells (Fig 1b, unlabeled). As expected^15^, optogenetic activation (480nm) of ChRoME-expressing presynaptic cells produced a sustained step-like current consistent with direct channel opening by light (Fig 1c). Neighboring cells expressing neither the ChRoME optogenetic tool nor the HyCal receptor did not exhibit light-activated currents (Fig 1d). But when we optogenetically stimulated the presynaptic neurons, we observed distinct postsynaptic currents in co-cultured cells expressing the postsynaptic HyCal receptor (Fig 1e). These results indicate the existence of synthetic synaptic connections between the presynaptic HyPep- and postsynaptic HyCal-expressing cells. In light of the absence of currents without the receptor (Fig 1d), we conclude that these results show the creation of a *de novo* synapse specifically through the reconstitution of the HySyn system.

We observed that repeated optogenetic stimulation of the presynaptic HyPep-expressing cells produced an integrating current in the postsynaptic HyCal receptor expressing cells (Fig 1h). With each stimulation, the inward current (downward trace deflections) increased to a new plateau level. This behavior is consistent with the known properties of both the receptor and neuromodulation by peptides. Specifically, the HyCal receptor does not desensitize^6^, and neuropeptides act at low concentrations across large volumes^8^. Thus, this integrating current suggests that an increasing fraction of HyCal channels open as neuropeptide release and stimulation persist. The selective presence of these currents in HyCal-expressing postsynaptic cells aligns with the expected specificity of this system. These data show that combining presynaptic HyPep and postsynaptic HyCal creates an artificial coupling of activity in a vertebrate neuronally-derived cell culture model, likely by utilizing volume transmission.

Next we used calcium imaging to examine the extent and reliability of HySyn neuromodulation in a population of cells. We used our established approach to activate presynaptic HyPep-expressing cells through optogenetics, but used the red-shifted variant Chrimson^16^ (Fig 2a, magenta cells) that is compatible with GCaMP imaging. In parallel, we monitored activation of postsynaptic cells with the calcium-sensitive fluorophore GCaMP6f^17^ (Fig 2a, green cells). In GCaMP-labeled cells lacking the HyCal receptor, we did not observe light-evoked calcium activity (Fig 2b), which was consistent with our electrophysiological data (Fig 1d). These results indicate that HyPep is inert to mouse Neuro2A cells. In contrast, when ‘presynaptic’ HyPep-positive cells were optogenetically stimulated, co-cultured cells expressing the postsynaptic HyCal receptor showed rising GCaMP signals (Fig 2c). In Figure 2c, responses are organized based on the response magnitude to highlight the frequency of responses. We classified approximately 34% (14/44) of quantified cells as clear ‘responders’ based on a change in GCaMP signaling greater than 3x standard deviations above those found prior to light stimulation. Quantification of this increase in calcium over the course of the 4-min stimulation (Figure 2d) highlights an overall doubling of the calcium signal compared to the initial signal, but some cells experienced changes in the GCaMP signal as high as 6-fold. We did not find a clear spatial pattern of activation with respect to the location of the presynaptic cells. This observation is consistent with our electrophysiological data and our expectations for a neuropeptide that functions through volume transmission. Postsynaptic calcium responses selectively in HyCal-expressing cells provides further support for the reconstitution of the HySyn synapse and the specificity of HySyn. Our finding that robust calcium response occur in a third of Neuro2a cells is consistent with a neuromodulatory function. Together, these observations support the idea that HySyn robustly creates a neuromodulatory synapse in a vertebrate neuronally-derived cell culture model. Our methods also demonstrate that HySyn is compatible with existing optogenetic and calcium imaging technologies.

**Figure 2.**
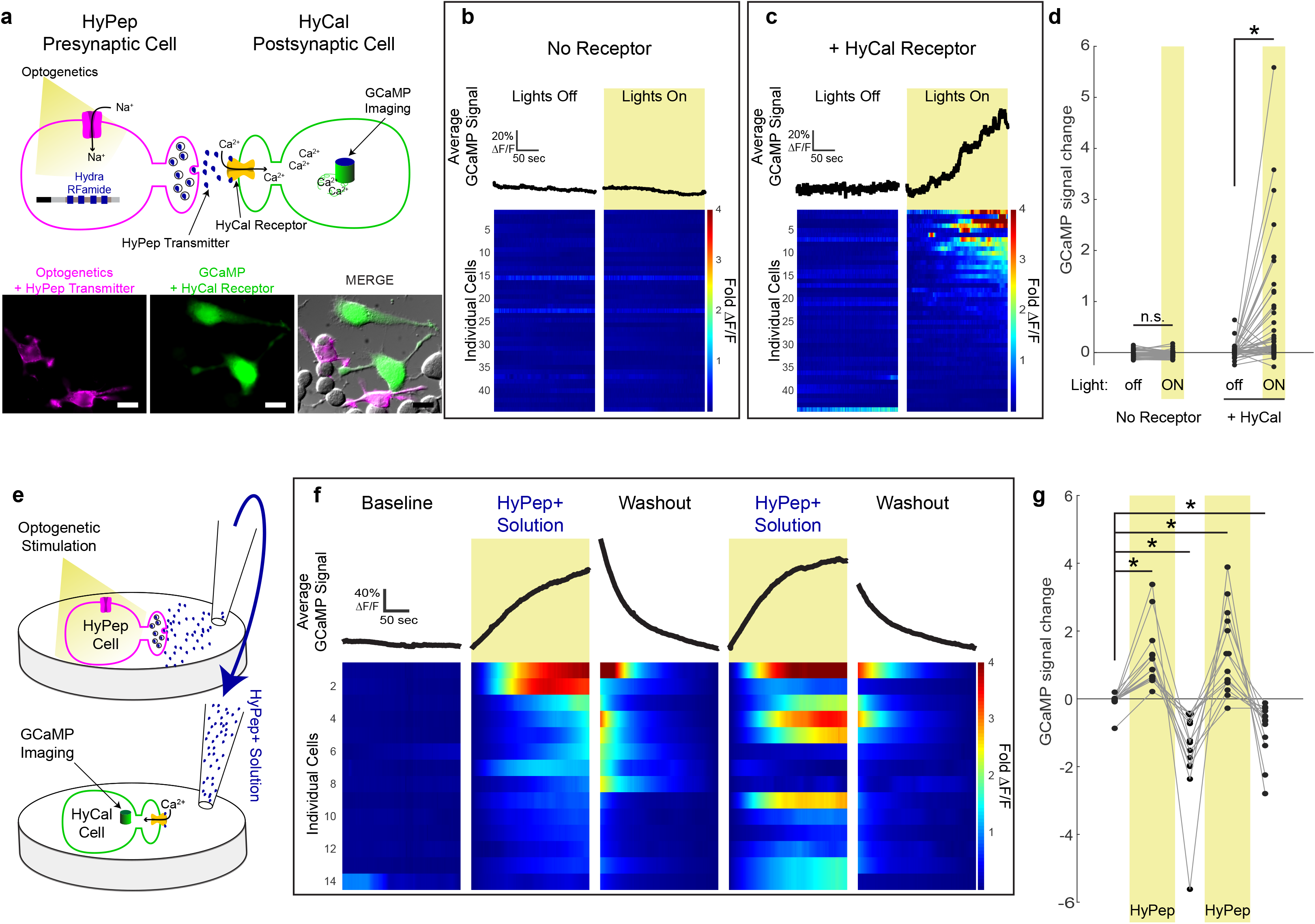
HySyn produces volumetric neuromodulation of calcium. **a**, Schematic illustrating the experimental paradigm used to characterize HyPep to HyCal synaptic communication with calcium imaging, top. Micrograph illustrating transfected Neuro2a cells used in these experiments, bottom (20μm scale bar). Chrimson was used for optogenetic stimulation (591nm, 500ms at 1Hz) of presynaptic cells expressing the HyPep carrier of the neuropeptide (‘Hydra RFamide’). GCaMP was expressed in distinct cells either alone (**b**) or with the cognate receptor HyCal (**c**). **b**, Average GCaMP signals, top, and individual cell heatmap profiles, bottom, for Neuro2a cells expressing GCaMP alone. Without receptor expression, GCaMP signals remained stable both without (left, ‘Lights Off’) and with (right, ‘Lights On’, yellow shading) optogenetic stimulation. **c**, GCaMP signals (as in **b**) except in cells also expressing the HyCal receptor illustrates the observed calcium signal rise over the course of light stimulation. Out of 44 cells, ~34% (14) showed changes in fluorescence over 3 standard deviations beyond the mean change observed prior to light stimulation. **d**, Quantification of the change in GCaMP signal (Final ΔF/F - Initial ΔF/F) revealed no effect of optogenetic stimulation by expressing GCaMP alone, but a doubling of intracellular calcium signal in postsynaptic cells expressing the HyCal receptor. **e**, Schematic illustration of solution transfer experiments. After optogenetic stimulation (as in **a** for 5min), bathing solution from HyPep-expressing cells (‘HyPep+ Solution’) was transferred to naive postsynaptic cells in another culture dish expressing GCaMP and the HyCal receptor. **f**, Following a stable baseline, applying the HyPep+ solution increased the GCaMP signal and this rise was reversed after washout and applying fresh bathing solution (‘Washout’). Repeated cycles of washout and application of the HyPep+ solution reproducibly increased the GCaMP signal in the same responding cells. **g**, Quantification (as in **d**) illustrates a doubling of signal after adding HyPep+ and halving of signal upon removal. * indicate p<0.05 using Mann-Whitney-Wilcoxon test.

We reasoned that if HySyn acts through volume transmission, optogenetic stimulation of a culture of ‘presynaptic’ HyPep-expressing Neuro2a cells would generate a HyPep+ solution that could act as a chemogenetic stimulator when added to a separate culture of ‘postsynaptic’ HyCal-expressing cells (Fig 2e). To test this idea, we collected the solution from optogenetically stimulated HyPep-expressing Neuro2a cells (HyPep+ solution) and added it to a separate HyCal-expressing cell culture. Prior to adding media, GCaMP signals were stable in all HyCal-expressing postsynaptic Neuro2A cells (Fig 2f), including the 14 cells (~45% of 31 examined) that ultimately showed responses to at least one application of HyPep+ solution. Application of HyPep+ solution produced an increase in the calcium signal, while subsequent washout and application of untreated solution led to a decline in the calcium signal (quantified in Fig 2g). When the Hypep+ solution was repeatedly added after washout cycles, and individual responding cells were tracked, we observed that of the responding Neuro2a cells, the majority (8/14) responded during both HyPep+ solution applications with a calcium rise greater than 3 standard deviations beyond any changes observed before application. We note however that cells responded differentially to the two applications, and that some cells responded to only a single HyPep+ solution application. These results are consistent with the expected neuromodulatory nature of the HySyn system, and suggest that HyCal interacts with cellular physiology to increase responsiveness of Neuro2a cells, but that it is not the sole determinant of activity. Importantly, these results provide further support that HySyn functions through volume transmission and that the HyCal receptor may be used with exogenously applied neuropeptide for pharmacological or chemogenetic manipulations of intracellular calcium.

We then sought to determine whether HySyn can modulate organismal behavior *in vivo*. In the nematode *C. elegans*, we expressed GFP tagged-HyPep throughout the nervous system (Fig 3a) and genetically targeted the HyCal receptor to muscle tissue. We predicted that activating the HyCal receptor in muscles through the release of HyPep from neurons would produce a spastic paralysis that would prevent proper worm movement. Consistent with our hypothesis, we observed that the engineered nematodes with neuro HyPep and muscle HyCal were severely uncoordinated, with signs of spastic paralysis by excessive muscular contraction (Supplementary Video S1). Transgenic animals performed very little crawling behavior, as illustrated by the trajectory of worms observed for 30min on an agar pad (Fig 3b, right panels). This behavioral outcome manifested in short trajectories and a reduced velocity of travel (Fig 3c, blue). *In vivo* expression of the pan-neuronal HyPep alone did not produce any observable phenotype or locomotor effect (Fig 3c, green), consistent with the specificity of the neuropeptide. Similarly, expression of the HyCal receptor did not produce locomotor defects by itself, and HyCal expression in muscle with HyPep expression in the intestines was insufficient to drive paralysis (Fig 3c, ‘Intestinal HyPep + Muscle HyCal’, magenta).

**Figure 3.**
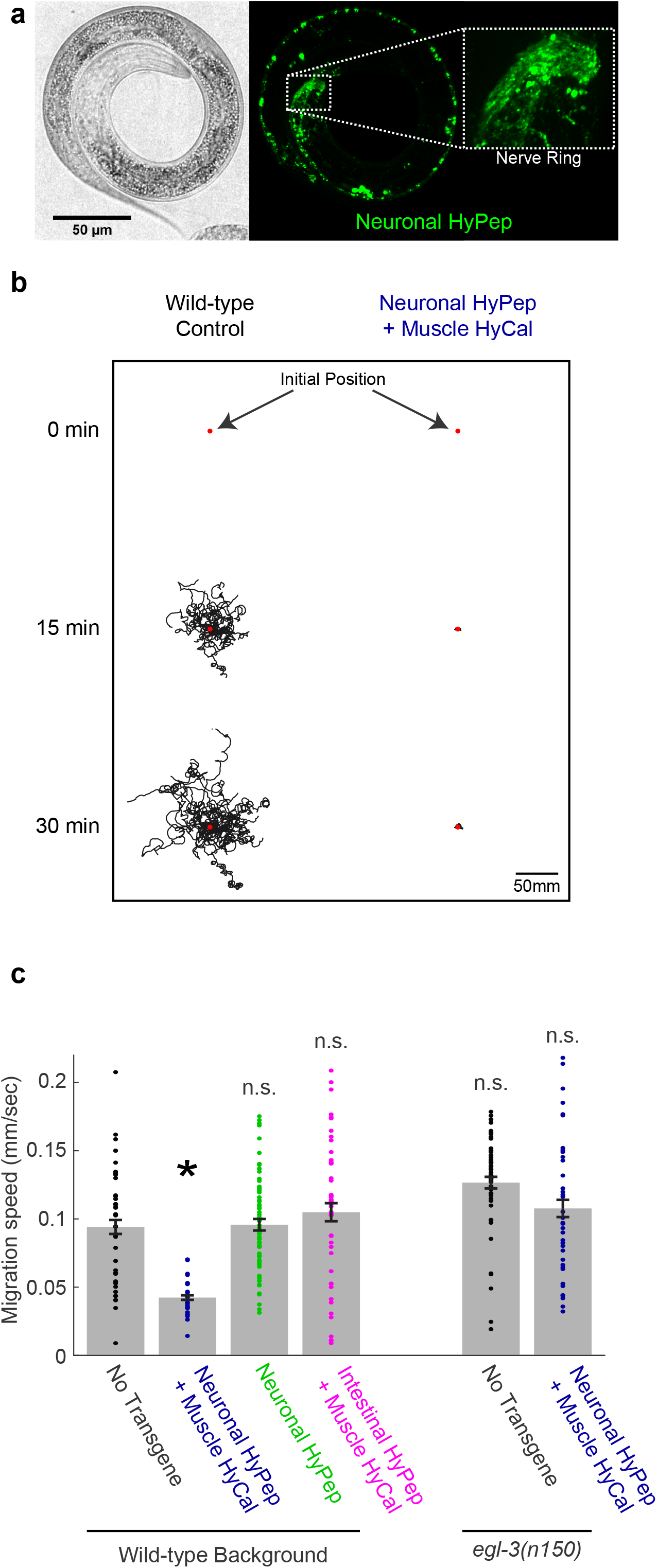
HySyn can alter organismal behavior. **a**, Bright-field (left) and fluorescence (right) micrographs show the pattern of expression for HyPep-GFP under the control of a neuronal promoter in the nematode *C. elegans*. The fluorescence pattern (green, right panel) shows a dense collection of puncta in the nerve ring (inset), a synaptically rich neuropil. **b**, Illustration of trajectories for 24 worms over a 30-min observation period. Each line represents a single worm track from a commonly-aligned initial position (red dot) after either a 0-min (top), 15-min (middle) or 30-min (bottom) monitoring period. Compared to the dispersion of wild-type control (left), transgenic animals expressing the synthetic HySyn synapse between neurons and muscles (right, ‘Neuronal HyPep + Muscle HyCal’) showed substantially reduced migration over time. **c**, During infrequent bouts of detectable migration in this 30-min interval, those expressing the full HySyn system (‘Neuronal HyPep + Muscle HyCal’, blue) move at a slower speed than control animals without HySyn (‘No Transgene’, black). Neither the neuropeptide itself (‘Neuronal HyPep’, green) nor the receptor in the presence of intestine-produced neuropeptide (‘Intestinal HyPep + Muscle HyCal’, magenta) altered migration speed. Migration in animals with a mutation in an essential and endogenous neuropeptide processing gene (‘*egl-3(n150*’) suppressed the function of the reconstituted HyPep-HyCal synapse (right-most bar, blue, compared to wild-type or *egl-3(n150)* mutant animals without transgene expression). * indicate p<0.05 using Mann-Whitney-Wilcoxon test.

The HyPep carrier initially exists as a pre-pro-peptide that requires cleavage by an endogenous pre-pro-convertase to create the final active peptide. This requirement provided an opportunity to use genetics to examine the specificity of our system *in vivo*. Specifically a mutant for the endogenous pre-pro-convertase enzyme in *C. elegans* predicted to cleave and produce the active peptide, *egl-3*. We generated animals containing the functional HySyn system, but in an *egl-3(n150)* mutant background, which lack a functional endogenous pre-pro-convertase (Fig 2c). Although the HySyn system components were both present, the ability of HySyn to produce paralysis was suppressed in the *egl-3(n150)* mutants, and worm locomotion reverted to wild type levels (Fig 3c, right-most bar). These results support the specificity of this system by demonstrating that functional presynaptic HyPep is required to produce the observed paralysis phenotype with postsynaptic HyCal. This outcome also highlights the potential of this system for forward genetic screening to identify novel components of neuropeptide processing and/or release, which can rescue this paralysis. Importantly, these results show that the HySyn system can reconfigure neural circuits *in vivo* to alter organismal behavior or probe genetic requirements of neuropeptide signaling and neuromodulation.

The versatility of our engineered HySyn system is illustrated by functioning in both mammalian neuronal cells and *in vivo C. elegans*. The system is also modular, which facilitates modifying and integrating its components with other approaches. For example, the GFP-labeled version of HyPep can be used on its own to track neuropeptide processing, trafficking or release. Also, the *Hydra* RFamide can be replaced with a single endogenous neuropeptide sequence to isolate function within complex neuropeptide precursors. On the other hand, the HyCal receptor can be utilized in chemogenetic approaches to pharmacologically enhance calcium in genetically targeted cells.

The ability to reconstitute genetically targetable, specific and functional synthetic synapses enables hypothesis-driven examination of sufficiency experiments for synaptic connections in circuit function and behavior. This, in turn, facilitates dissecting the behavioral role of specific neural circuit configurations. The power of the HySyn system lies in the ability to create a new functional synaptic connection to bias the relationship between a particular synaptic input and postsynaptic intracellular calcium. Calcium, in turn, can alter synaptic plasticity, gap junction function, and gene expression at different timescales and based on the persistence of the signal^16^. The HySyn system is fully compatible with existing tools to optogenetically or chemogenetically control neural circuits, but provides an innovative and complementary avenue to control the wiring of these circuits. As a volume-transmission neuromodulatory connection, this synthetic synapse can be applied in a wide range of neuroanatomical configurations, including in the absence of direct ultrastructural synaptic contact. Our engineered HySyn system enables biasing or reconfiguring neural circuits into discrete states for *in vivo* dissection of the role of neuromodulation to establish neural circuit logic and connectivity.

## Acknowledgements

This work was mainly conducted in the Grass Laboratory at the Marine Biological Laboratories (MBL) with funding through a Grass Fellowship awarded to J.D.H. Thanks to Richard Goodman (OHSU) for encouragement during conceptualization of the fellowship application, and the 2019 Grass Fellows, Mel Coleman (Grass Director) and Christophe Dupré (Associate Director) for advice and support during the summer fellowship. We thank the MBL Division of Education and participants in the Vendor Equipment Loan Program. Special thanks to Sutter Instruments, who generously provided all electrophysiology equipment and substantial on-site assistance, and Zeiss, who provide on-site assistance at MBL. We thank Zhao-Wen Wang and Ping Liu (UConn) for guidance and training in patch-clamp electrophysiology, as well as providing Neuro2a cells. We thank Rob Steele (UCI) for supplying *Hydra*, as well as advice and inspiration on *Hydra* biology. We thank members of the Colón-Ramos lab and Hari Shroff (NIH) for thoughtful comments on the manuscript, and Life Science Editors for editing assistance. D.A.C.-R. is an MBL Fellow. Research in the D.A.C.-R. lab was supported by NIH R01NS076558, DP1NS111778 and by an HHMI Scholar Award.

## Author Contributions

J.D.H designed, performed and analyzed experiments. J.D.H. and D.A.C.-R. prepared the manuscript.

## Materials and Methods

### Molecular Biology

All HySyn synaptic components were synthesized (Gene Universal, Newark DE USA) with flanking attB1/B2 sites for subsequent Invitrogen BP Gateway recombinational cloning into pDONR221 entry vector (Thermofisher, Waltham MA USA). To produce mammalian expression constructs, Gateway LR recombination was performed into the pEZY3 expression construct (AddGene 18672). For *C. elegans* expression, LR recombination subcloning with a Multisite Gateway system^18^ was used to generate the presynaptic HyPep construct under the control of the pan-neuronal rab-3 promoter or the postsynaptic HyCal receptor under the control of the muscle-specific myo-3 promoter. Optogenetic and calcium imaging plasmids were obtained from Addgene (ChRoME^15^, 108902; ChRimson^15^, 105447; GCaMP6f^17^, 40755). Detailed cloning information will be supplied upon request.

### Cell Culture & transfection

Neuro2a neuroblastoma cells (gift from Zhao-Wen Wang, UConn) were cultured in Opti-MEM (Thermofisher) supplemented with 5% fetal bovine serum (FBS, Thermofisher) and penicillin/streptomycin. Cell transfection was performed with Lipofectamine 2000 (Thermofisher) in Opti-MEM media according to the manufacturer’s protocol. Transfections were performed in separate dishes for the pre- and post-synaptic components. After ~24hr, transfected cells were dissociated with trypsin-EDTA (Thermofisher) followed by co-culture of separately transfected populations on poly-L lysine coated coverslips. Electrophysiology and calcium imaging were performed 24-48hr after co-culture.

### Electrophysiology and calcium imaging

Coverslips with Neuro2a cells were mounted in the QE-1 chamber (Warner Instruments, Hamden CT USA) on an MT1000 stage (Sutter Instruments, Novato CA USA) under an Axio Vert.A1 microscope (Zeiss, Jenna, Germany). Cells were bathed in an external solution containing (in mM): 140 NaCl, 1.3 KCl, 4 CsCl, 2 TEACl, 1 NaH_2_PO_4_, 1.8 CaCl, 0.8 MgSO_4_, 5.5 D-glucose, 10 HEPES. Illumination for imaging and optogenetics was achieved using the Lambda-421 optical beam combiner (Sutter Instruments). Patch pipettes were pulled with a P1000 micropipette puller (Sutter Instruments) to 3-5 MΩ, then filled with internal solution containing (in mM): 140 KCl, 1 MgCl_2_, 5 K_4_BAPTA, 3 CaCl_2_, 25 HEPES. Electrophysiological data was acquired with the Double Integrated Patch Amplifier and Data Acquisition unit and SutterPatch software (Sutter Instruments). Solution transfer and washout experiments for calcium imaging were achieved manually with the aid of a micropipette in a total 1ml bath solution. Calcium imaging, also on the Axio Vert.A1, was captured with a ORCA-Flash4.0 LT (Hamamatsu, Hamamatsu City Japan) controlled by μManager^3^. Image quantification and planned comparisons using non-parametric Mann-Whitney-Wilcoxon test were implemented in Matlab.

### Worm transgenesis and behavior

Transgenic lines were created by microinjection into the distal gonad syncytium as previously described^4^ and selected based on expression of co-injection markers, Punc-122∴GFP or Punc-122∴dsRed. Confocal images of transgene expression were acquired using Volocity (Perkin Elmer) on the UltraView VoX spinning disc confocal microscope with a NikonTi-E stand and a 60X CFI Plan Apo VC, NA1.4, oil objective. Figures were prepared with FIJI^19^ and Adobe Illustrator. Animal migration on an agar pad was monitored for 30 min at 2fps using a MightEx camera (BCE-B050-U). Trajectories were analyzed using an adaptation of the MagatAnalyzer software package as previously described^20^. Track analyses and planned comparisons using non-parametric Mann-Whitney-Wilcoxon test were implemented in Matlab.

---

**Figure S1.**
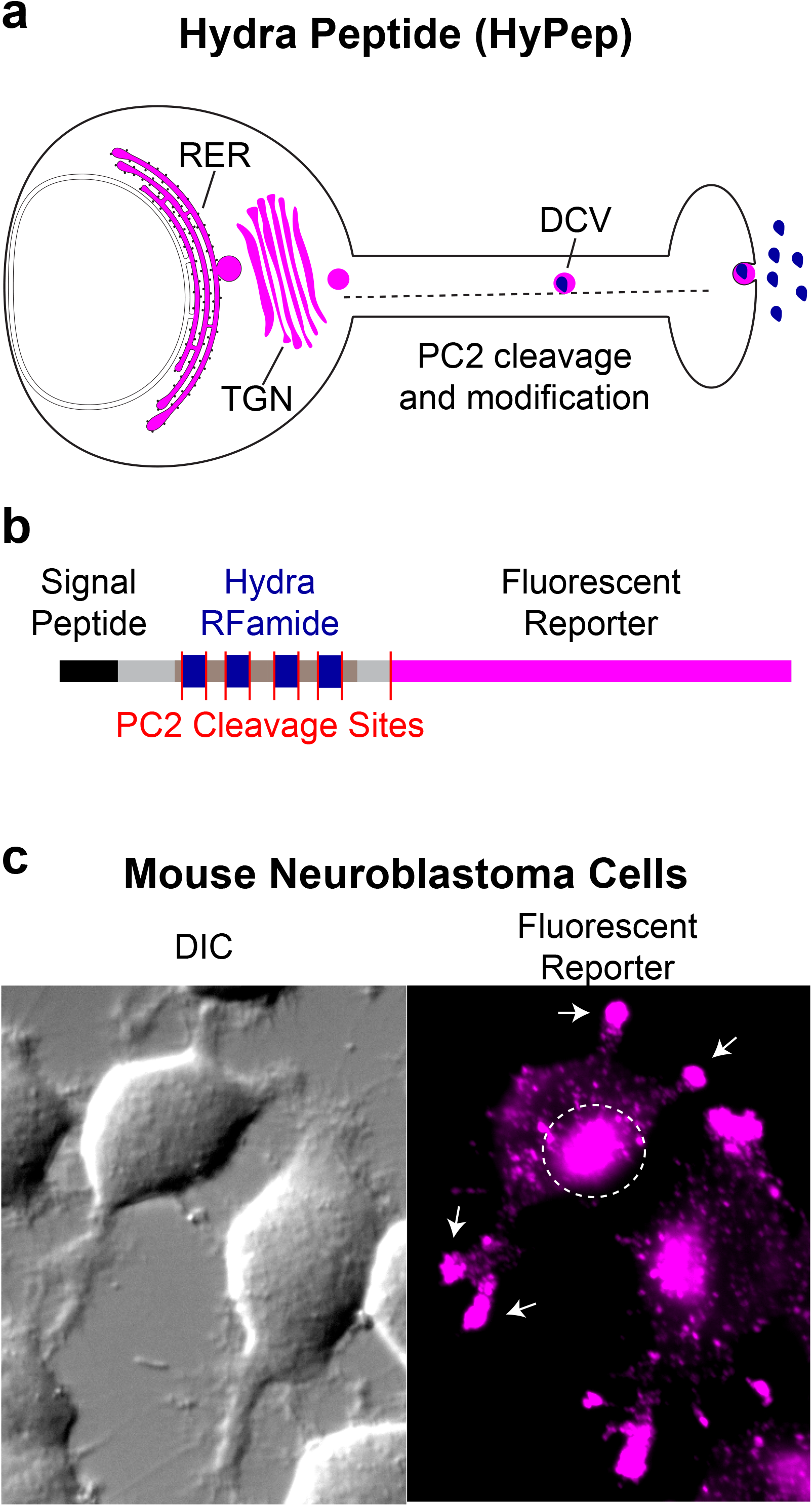
*Hydra*-derived neuropeptide (HyPep) carrier for heterologous expression, processing and transport of neuropeptide. **a**, Illustration of the cellular neuropeptide processing pathway utilized by the *Hydra*-derived neuropeptide (HyPep). The regulated exocytosis pathway involves synthesis in the rough endoplasmic reticulum (RER), trafficking through the trans-golgi network (TGN), and packaging into dense core vesicles (DCVs) for release. **b**, Schematic of the HyPep pre-pro-peptide carrier molecule for peptide processing and delivery. An N-terminal signal peptide (‘Signal Peptide’), based on the broadly expressed Neuropeptide Y sequence, targets the precursor molecule to the RER for synthesis. Cleavage at consensus recognition sites (red lines) for the endopeptidase pre-pro-convertase 2 (PC2) produces individual peptide fragments for further chemical modification into active neuropeptides (‘Hydra RFamide’). **c**, Expression of the HyPep-GFP reporter in N2A neuroblastoma cells reveals localization in a pattern consistent with sites of neuropeptide synthesis and trafficking, including expression in compartments appearing to be perinuclear (as expected for the RER, white dashed circle) and at neurite extensions (white arrows).

**Supplementary video S1.** Swimming behavior in transgenic *C. elegans* expressing the synthetic neuron-to-muscle HySyn synapse. Order of display: Wild-type *C. elegans*, Neuro-Muscular HySyn, *egl-3(n150)* animals, Neuro-Muscular HySyn in *egl-3(n-150)* background.

## References

1. Roth, B. L. DREADDs for Neuroscientists. Neuron 89, 683–694 (2016).

2. Kim, C. K., Adhikari, A. & Deisseroth, K. Integration of optogenetics with complementary methodologies in systems neuroscience. Nat. Rev. Neurosci. 18, 222–235 (2017).

3. Bargmann, C. I. & Marder, E. From the connectome to brain function. Nat. Methods 10, 483–490 (2013).

4. Nadim, F. & Bucher, D. Neuromodulation of Neurons and Synapses. Curr. Opin. Neurobiol. 0, 48–56 (2014).

5. Assmann, M., Kuhn, A., Dürrnagel, S., Holstein, T. W. & Gründer, S. The comprehensive analysis of DEG/ENaC subunits in Hydra reveals a large variety of peptide-gated channels, potentially involved in neuromuscular transmission. BMC Biol. 12, 84 (2014).

6. Dürrnagel, S., Falkenburger, B. H. & Gründer, S. High Ca(2+) permeability of a peptide-gated DEG/ENaC from Hydra. J. Gen. Physiol. 140, 391–402 (2012).

7. Findeisen, M., Rathmann, D. & Beck-Sickinger, A. G. RFamide Peptides: Structure, Function, Mechanisms and Pharmaceutical Potential. Pharmaceuticals 4, 1248–1280 (2011).

8. Li, C. & Kim, K. Neuropeptides. WormBook Online Rev. C Elegans Biol. 1–36 (2008) doi:10.1895/wormbook.1.142.1.

9. Carafoli, E. & Krebs, J. Why Calcium? How Calcium Became the Best Communicator. J. Biol. Chem. 291, 20849–20857 (2016).

10. Wei, S., Cassara, C., Lin, X. & Veenstra, R. D. Calcium–calmodulin gating of a pH-insensitive isoform of connexin43 gap junctions. Biochem. J. 476, 1137–1148 (2019).

11. Burke, K. J., Keeshen, C. M. & Bender, K. J. Two Forms of Synaptic Depression Produced by Differential Neuromodulation of Presynaptic Calcium Channels. Neuron 99, 969–984.e7 (2018).

12. Whim, M. D. & Moss, G. W. A novel technique that measures peptide secretion on a millisecond timescale reveals rapid changes in release. Neuron 30, 37–50 (2001).

13. Han, W., Ng, Y.-K., Axelrod, D. & Levitan, E. S. Neuropeptide release by efficient recruitment of diffusing cytoplasmic secretory vesicles. Proc. Natl. Acad. Sci. U. S. A. 96, 14577–14582 (1999).

14. Remacle, A. G. et al. Substrate Cleavage Analysis of Furin and Related Proprotein Convertases. J. Biol. Chem. 283, 20897–20906 (2008).

15. Mardinly, A. R. et al. Precise multimodal optical control of neural ensemble activity. Nat.Neurosci. 21, 881–893 (2018).

16. Klapoetke, N. C. et al. Independent optical excitation of distinct neural populations. Nat.Methods 11, 338–346 (2014).

17. Chen, T.-W. et al. Ultrasensitive fluorescent proteins for imaging neuronal activity. Nature 499, 295–300 (2013).

18. Hawk, J. D. et al. Integration of Plasticity Mechanisms within a Single Sensory Neuron of C. elegans Actuates a Memory. Neuron 97, 356–367.e4 (2018).

19. Schindelin, J. et al. Fiji: an open-source platform for biological-image analysis. Nat. Methods 9, 676–682 (2012).

20. Gershow, M. et al. Controlling airborne cues to study small animal navigation. Nat. Methods 9, 290–296 (2012).

